# Genome-wide CRISPR screen identifies AC9 as a key regulator of ER Ca^2+^ homeostasis involved in neuronal differentiation

**DOI:** 10.1101/2024.02.05.578803

**Authors:** Liuqing Wang, Jia Li, Yiping Wang, Ziyi Zhong, Yuqing Wang, Rui Huang, Bingwei Zhang, Panpan Liu, Erkejiang Ye, Ruotong Cao, Sher Ali, Yuepeng Ke, Junjie Yang, Tatsushi Yokoyama, Jin Liu, Xiaoyan Zhang, Masayuki Sakamoto, Lin Sun, Yubin Zhou, Youjun Wang

## Abstract

Endoplasmic reticulum (ER) calcium (Ca^2+^) homeostasis is essential for maintaining normal cellular physiological functions. Its disturbance is strongly linked to the onset and progression of human diseases, including cancer, developmental defects, and neurodegenerative disorders. The lack of sensitive ratiometric ER Ca^2+^ indicators, nevertheless, hinders systematic investigation of ER Ca^2+^ modulators and the underlying mechanisms. Capitalizing on two ultra-sensitive ER Ca^2+^ indicators and CRISPR-based genome-wide screening, we identified a set of proteins capable of reducing the ER Ca^2+^ content. Further comparative analysis and qPCR validation pinpointed adenylate cyclase 9 (AC9), which is upregulated during neuronal differentiation, as a key ER-Ca^2+^-reducing regulator. Mechanistically, AC9-mediated production of cAMP is not essential for its ability to reduce ER Ca^2+^ content. Instead, AC9 inhibits store operated calcium entry (SOCE) by acting on Orai1, ultimately causing attenuation of ER Ca^2+^ level. More physiologically relevant, upregulation of AC9 in neurons is essential for reducing ER Ca^2+^ levels during *Drosophila* brain development. Collectively, this study lays a solid groundwork for further in-depth exploration of the regulatory mechanisms dictating ER Ca^2+^ homeostasis during neuronal differentiation and brain development.

## Introduction

Calcium (Ca^2+^) ion serves as a universal secondary messenger that regulates a myriad of biological processes, spanning from cell division to apoptosis, and from early embryonic development to aging[1]. During early development, Ca^2+^ signals help establish the dorsal–ventral (D–V) axis and contribute to organogenesis[2]. In zebrafish embryos, injection of a Ca^2+^ buffer, 1,2-bis(o-aminophenoxy)ethane-N,N,N′,N′-tetraacetic acid (BAPTA), into zygotes results in zygotic cleavage failure and retains embryos at the one-cell stage[3]. Loading blastomeres with BAPTA has been shown to disrupt motor neuron development[4]. Endoplasmic reticulum (ER) is a major source of Ca^2+^ for generating cytoplasmic Ca^2+^ signals. Inhibition of sarco/endoplasmic reticulum Ca^2+^-ATPase (SERCA) by thapsigargin (TG) during gastrulation induces cyclopia[5] and affects left-right asymmetry of the heart and brain[6]. Beyond developmental processes, ER Ca^2+^ homeostasis is crucial for ensuring proper cellular activities. Dysregulation of ER Ca^2+^ levels can severely impair the functions of the cardiovascular, nervous and immune systems, causing pathological conditions like arrhythmia[7], immunodeficiency[8], and neurodegeneration[9]. Hence, restoring ER Ca^2+^ balance might represent a vital strategy for addressing these diseases.

Maintaining precise ER Ca^2+^ homeostasis relies on the intricate control over bidirectional transfer of Ca^2+^ across the ER membrane: both efflux and influx processes. Regarding efflux, the ER Ca^2+^ release channels, such as inositol 1, 4, 5-trisphosphate receptors (IP_3_Rs) and ryanodine receptors (RYRs), are activated upon ligand binding, allowing Ca^2+^ to be released from the ER/SR into the cytoplasm. Additionally, studies have reported a protein called TMCO1 on the ER membrane, which may form a Ca^2+^ leak channel after ER Ca^2+^ overload, allowing Ca^2+^ to leak into the cytoplasm[10]. In terms of influx, SERCA transports Ca^2+^ back into the ER from the cytoplasm, replenishing the ER Ca^2+^ store and restoring Ca^2+^ levels to its resting state. Of note, a crucial mechanism for maintaining ER Ca^2+^ homeostasis is store-operated Ca^2+^ entry (SOCE)[11, 12]. SOCE is initiated by ER Ca^2+^ store depletion, which activates the ER-resident Ca^2+^ sensor, stromal interaction molecule 1 (STIM1), via Ca^2+^-depletion induced conformational changes and oligomerization. Activated STIM1 proteins subsequently translocate to the ER-plasma membrane (PM) junctions, where they directly engage and gate the PM-embedded Ca^2+^ channel Orai1 to trigger the influx of extracellular Ca^2+^ into the cytoplasm, thereby replenishing the depleted ER Ca^2+^ store and restoring intracellular Ca^2+^ homeostasis[13].

Multiple cellular factors modulate the activity of these Ca^2+^ handling machineries. Previous studies have found that tumor suppressors, such as promyelocytic leukemia protein (PML) and breast cancer 1 (BRCA1), enhance the function of IP_3_R, increasing IP_3_R-mediated Ca^2+^ release[14, 15]. Several studies have shown that the apoptotic factor Bcl-2 could reduce the ER Ca^2+^ content and IP_3_R-mediated Ca^2+^ release by decreasing SERCA2b activity, or by increasing Ca^2+^ leakage[16–20]. SOCE-associated regulatory factor (SARAF) inactivates SOCE to prevent excessive Ca^2+^ refilling[21]. Additionally, an isoform of the secretory pathway Ca^2+^ ATPase (SPCA2) was shown to elevate basal Ca^2+^ levels via its direct interaction with Orai1[22]. While these reports have provided insights into the regulation of ER Ca^2+^ homeostasis, many of them employ indirect ways to detect ER Ca^2+^ levels by using cytoplasmic indicators such as Fura-2. Even studies using ER indicators like Cameleon or D1ER are limited by their non-optimal Ca^2+^ affinities for reporting ER Ca^2+^ signaling and low dynamic ranges. Due to the lack of ER indicators with high sensitivity and dynamics, a systematic exploration of protein modulators of ER Ca^2+^ homeostasis is currently hindered.

Utilizing two highly sensitive ER Ca^2+^ indicators recently developed by us, miGer[23] and TuNer-s[24], along with genome-wide CRISPR/Cas9 screening, we conducted a systemic search for modulators of ER Ca^2+^ homeostasis. We identified adenylyl cyclase 9 (AC9) as a key protein upregulated in the brain of fruit flies and rodents that account for reduced ER Ca^2+^ content during neuronal differentiation and brain development. Mechanistically, AC9 exerts its ER Ca^2+^-modulating effect by suppressing Orai1-mediated SOCE, a function that is independent on its enzymatic activity to convert ATP into cAMP. Together, our study establishes AC9 as a previously unrecognized regulator of ER Ca^2+^ homeostasis that might regulate neuronal development. Moreover, other ER Ca^2+^-modulating candidates emerged from our screening also form a solid foundation for future exploration of their functions.

## Results and Discussions

### Genome-wide CRISPR/Cas9 together with Ca^2+^-imaging-based screening identified genes lowering ER Ca^2+^ levels

To comprehensively explore key regulators of ER Ca^2+^ homeostasis, we employed a genome-wide CRISPR/Cas9 editing approach combined with a sorting method based on ratiometric fluorescence intensities indicative of ER Ca^2+^ levels (**Fig. 1A**). Firstly, we generated a HEK 293 stable cell line co-expressing Cas9 and the ratiometric ER indicator miGer (**Fig. 1B**). miGer comprises G-CEPIA1er a Ca^2+^ sensor, and mScarlet, a red fluorescence protein used as an expression marker. The fluorescence ratio (F_G-CEPIA1er_/F_mKate_) of miGer as a direct index of ER Ca^2+^ level. We transduced the GeCKOv2 library containing 123,411 sgRNAs targeting 19,050 annotated genes[25] to the stable cells, using a multiplicity of infection (MOI) of 5 to ensure only one sgRNA transfected per cell. Following a five-day selection with puromycin, the obtained cells were sorted by fluorescence-activated cell sorting (FACS) based on altered miGer ratios. We divided cells into different bins based on miGer ratios (control; ratio-high; ratio-low). DNA of these groups was extracted for next-generation sequencing (NGS). After filtering out irrelevant entries sgRNAs targeting 2,216 genes and microRNA were significantly enriched in the ratio-high group, and 1,381 in the ratio-low group.

**Figure 1.**
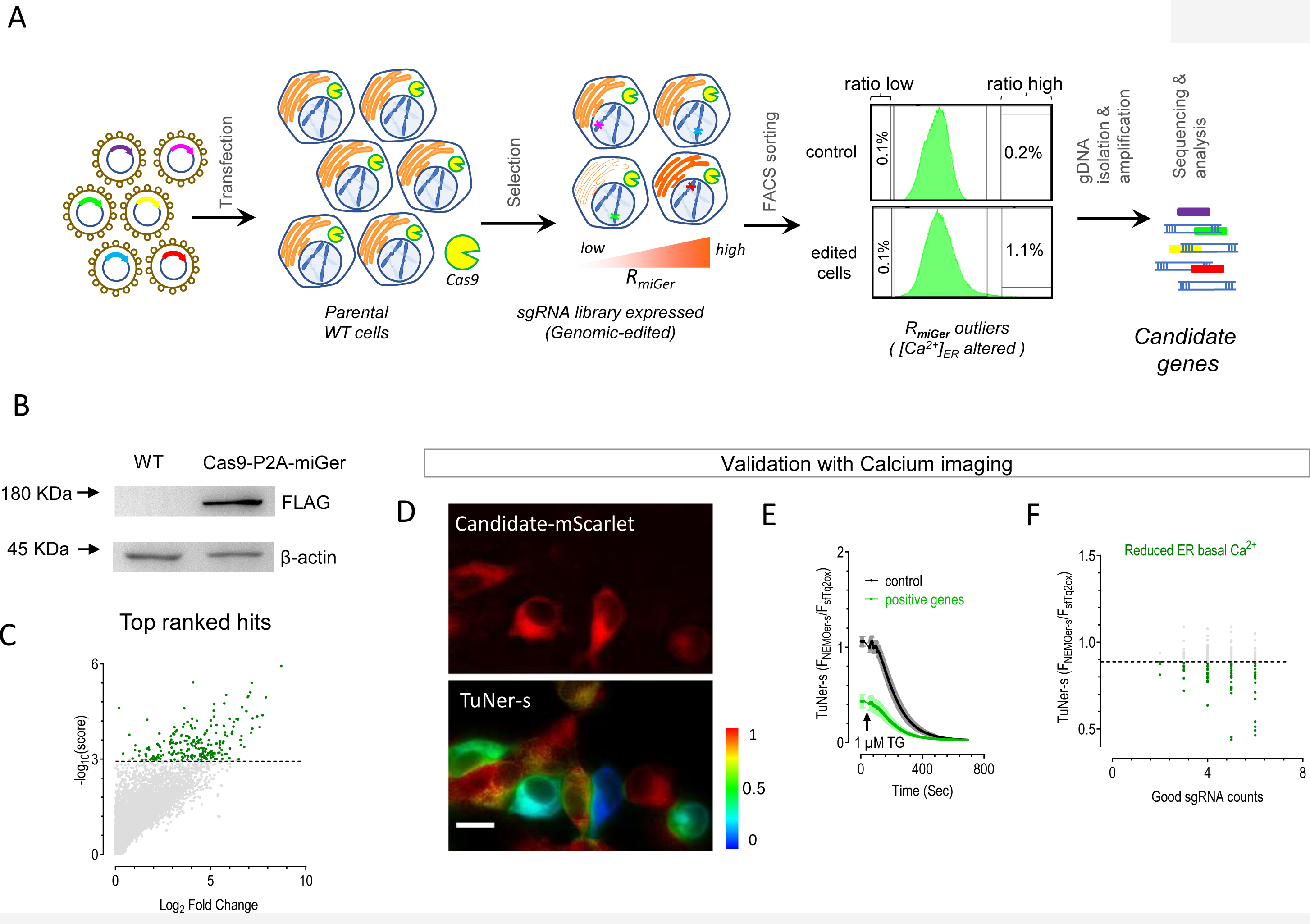
Regulators of ER Ca^2+^ levels unveiled through genome-wide CRISPR/Cas9 combined with Ca^2+^ imaging-based screening. A) Schematic diagram illustrating the workflow of genome-wide CRISPR/Cas9 screening for regulators of ER Ca^2+^ homeostasis. B) Western blot showing the expression of Flag-Cas9 in Cas9-P2A-miGer stable cells. C) Enriched genes identified from CRISPR/cas9-edited cells with increased ER Ca^2+^ levels. Top 1%-ranked genes by applying the robust ranking aggregation (RRA) algorithm are indicated in green. D-E) Imaging assays used to validate top-ranked hits using HEK293 cells stably expressing an ER Ca^2+^ sensor TuNer-s (TuNer-s cells). ER Ca^2+^ level was indicated by the ratio of TuNer-s (F_NEMOer-s_/F_sfTq2ox_). After acquiring basal images, ER Ca^2+^ stores were depleted with 1 μM Thapsigargin (TG). TuNer-s ratios of cells with mScarlet fluorescence (the experimental group) were compared with cells without mScarlet fluorescence (internal control) in the same view field. The TuNer-s ratios were normalized to those of the internal control cells. Unless specified, all imaging experiments in this work were performed and presented the same way. Representative basal fluorescence images under the same view field (D). Top, mScarlet fluorescence. Bottom, TuNer-s ratio. Scale bar, 10 μm. Typical traces (E). F) Summary plot showing effects of overexpressed top ranked hits on basal ER Ca^2+^ levels. Green dots represent validated genes via the aforementioned imaging assays shown in (D-E). While gray dots represent genes that showed no appreciable effects. Data are shown as the mean of at least three independent experiments.

We then conducted Robust Ranking Aggregation (RRA) analysis and selected the top 200 genes from each group for further verification with Ca^2+^ imaging (**Fig. 1C**). We expressed mScarlet-tagged candidates into HEK293 cells stably expressing a more sensitive Ca^2+^ indicator TuNer-s, which is constituted with NEMOer-s as a Ca^2+^ sensor and sfTq2ox acting as an expression reporter[24]. We further examined the effects of overexpressing candidates on the ER Ca^2+^ level indicated by TuNer-s ratio (F_NEMOer-s_/F_sfTq2ox_). mScarlet-negative cells in the same field were used as internal controls to eliminate possible artifacts (**Fig. 1D**). After collecting the basal ratios, we used 1 μM TG to fully deplete the ER Ca^2+^ store and ensured that the ratio in all cells dropped to the same level, confirming that the differences in resting values were due to varying ER Ca^2+^ levels and not affected by other factors. Candidates were considered positive hits based on their effects on ER Ca^2+^ levels. Those from the ratio-high group, showing significantly reduced TuNer-s ratios indicative of lower ER Ca^2+^ levels, as well as candidates in the ratio-low group that significantly increased ER Ca^2+^ levels, met the positive hit criteria (**Fig. 1E**). We did not obtain any positive hits from the ratio-low group. Even though this is consistent with results from FACS sorting showing that the cell proportion of the ratio-low group was similar to that of the non-edited group (**Fig. 1A**), it is interesting that not a single candidate gene from ratio-low group could significantly alter ER Ca^2+^ levels. Follow-on studies are needed to elucidate this in the future. However, among the 169 candidates in the ratio-high group that could be constructed into plasmids, 66 were found to significantly reduce resting ER Ca^2+^ levels (**Fig. 1F**). With a positive rate of 39%, this finding demonstrates the effectiveness of our screening in identifying ER Ca^2+^ regulators in mammalian cells.

### AC9 identified as a strong hit that is upregulated during neuronal development

We then performed GO enrichment analysis on the 66 genes that could reduce ER Ca^2+^ levels. Positive regulation of cell development was found to be among the top ranked biological processes (**Fig. 2A**), hence motivating us to investigate how ER Ca^2+^ homeostasis undergoes changes during cell differentiation and development, particularly in neurons. To test this, we induced the differentiation of Neuro-2a (N2a) cells, a neuroblastoma cell line stably expressing the ER indicator miGer. By treating them with reduced serum concentration (0.1% FBS) in DMEM in the presence of 20 µM retinoic acid (RA)[26], N2a cells will transform into neuron-like structures with extended axons. We monitored changes in the miGer signals throughout this differentiation process. As differentiation progressed, the miGer ratiometric signals gradually declined, indicating a reduction in the ER Ca^2+^ level (**Fig. 2B**). By contrast, the control group without induction showed no overt changes in the miGer signals (**Fig. 2C**). To further confirm our results, we used our newly developed highly sensitive ratiometric indicator, TuNer-s[24], to monitor changes in ER Ca^2+^ levels before and after N2a cell differentiation. Compared to the undifferentiated control group, ER Ca^2+^ levels significantly declined following 3-day treatment of N2a cells with RA (**Fig. 2D**). These results show a progressive reduction in ER Ca^2+^ levels during N2a cell differentiation, suggesting a close correlation between alterations in ER Ca^2+^ homeostasis and neuronal differentiation.

**Figure 2.**
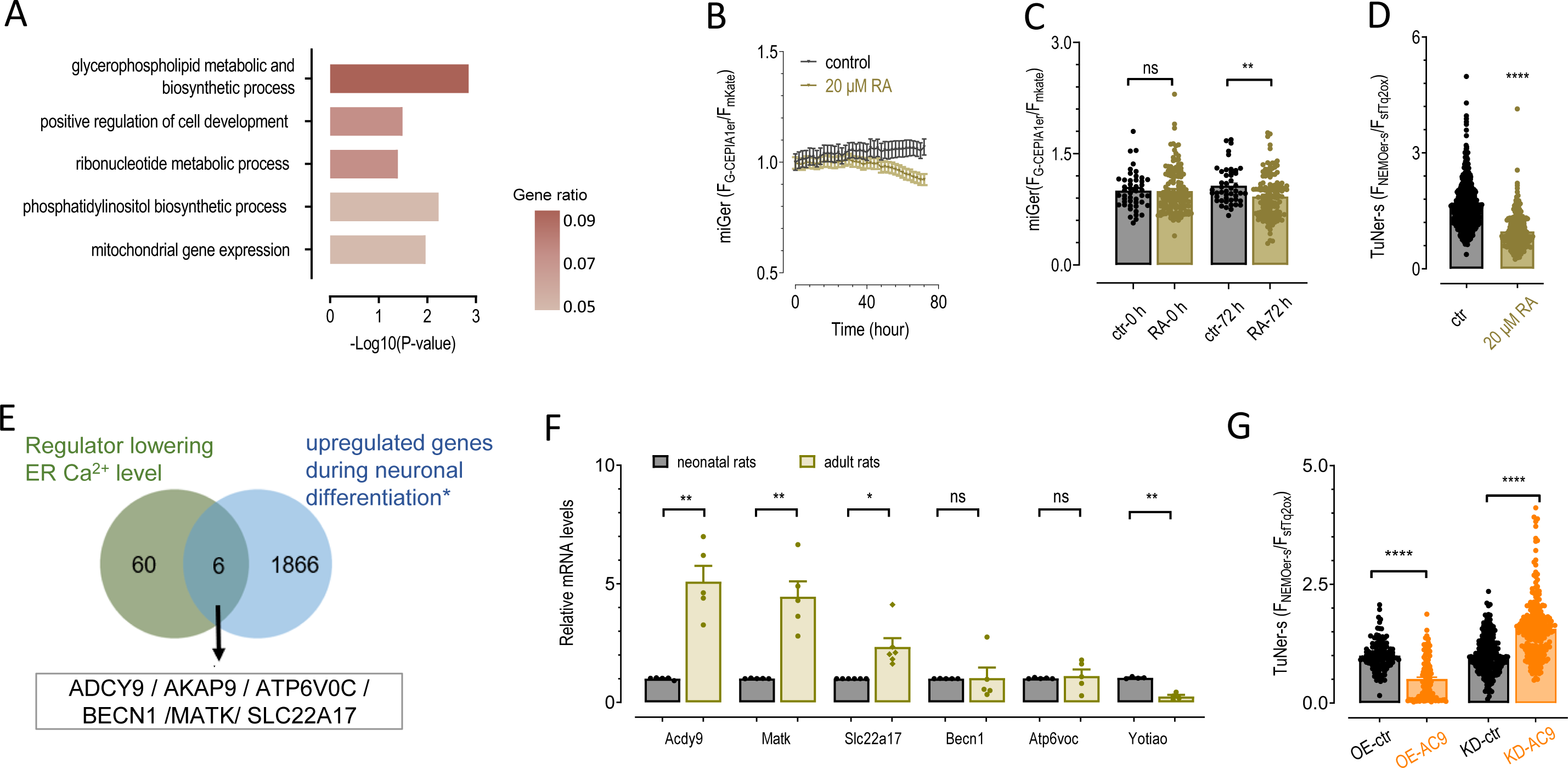
AC9 is a key regulator of ER Ca^2+^ homeostasis during neuronal differentiation and brain development. A) GO analysis of the identified genes that lead to reduced ER Ca^2+^ content. Top 5 enriched GO biological process terms are listed. B) Typical traces showing the changes of ER Ca^2+^ levels during differentiation of Neuro-2A (N2a) cells induced by 40 μM retinoic acid (RA). Control groups were treated with DMSO (n = 47 cells for control and n=134 cells for RA from three independent experiments). The ER-resident Ca^2+^ indicator miGer is used to report ER Ca^2+^ levels. The ratios of miGer were normalized to the ratio at the beginning of recording (time 0). C) Quantification of miGer signals at 0 and 72 h after RA treatments (\*\**P* < 0.01; ns, not significant; unpaired Student’s t-test). D) Statistics showing the effects of 20 μM RA on ER Ca^2+^ levels reported by a more sensitive ER Ca^2+^ indicator TuNer-s. The TuNer-s ratio is normalized to the control cells (n=773 cells for control and n= 209 cells for RA from at least three independent experiments. \*\*\*\**P* < 0.0001; unpaired Student’s t-test). E) Venn diagram showing the overlap between genes capable of reducing the ER Ca^2+^ level and those previously reported to be significantly upregulated (with *P*<0.05) during neuronal differentiation. F) Relative mRNA levels of overlapped genes in cerebral cortex from neonatal rat and adult rat. GAPDH was used as an internal control (Data are presented as the mean ±SEM of at least three independent experiments, each from the average of at least 3 technical replicates. \**P* < 0.05, ** *P* < 0.01; ns, not significant; Paired Student’s t-test). G) Effects of AC9 overexpression (OE) or knock-down (KD) on basal ER Ca^2+^ levels of N2a cells indicated by TuNer-s ratio (n=125 cells for OE-control, n=131 cells for OE, n=217 cells for KD-control and n=255 cells for KD. **** *P* < 0.0001, unpaired Student’s t-test). Data are shown as mean ± SEM of at least three independent experiments.

To explore factors contributing to the decline of ER Ca^2+^ levels during this differentiation process, we intersected the newly identified 66 positive hits with previously reported 1,872 gene that are differentially upregulated in neuronal cells during differentiation[27], which uncovered six overlapping genes, including AC9, AKAP9, ATP6V0C, MATK, BECN1, and SLC22A17 (**Fig. 2E**). Next, we asked whether the expression of these genes was indeed altered during development by examining their mRNA levels in rat cortical tissues. Compared to neonatal rats, the expression of *Adcy9* and *Matk* was significantly upregulated in the adult rat cortex, with AC9 showing a prominent 5-fold increase in expression (**Fig. 2F**). Given that AC9 exhibited the most notable upregulation, we selected AC9 for further investigation by either knocking down or over-expressing AC9 in N2a cells. AC9 overexpression in N2a-TuNer-s cells resulted in a significant decrease in ER Ca^2+^ levels, whereas knockdown of AC9 led to an appreciable increase in ER Ca^2+^ levels (Fig. 2G). Collectively, these results further validate the regulatory effects of AC9 on ER Ca^2+^ levels in neurons under physiological conditions.

### AC9 attenuates the ER Ca^2+^ level through a mechanism independent of its enzymatic function

AC9 is a membrane-localized adenylate cyclase that catalyzes the synthesis of cyclic AMP (cAMP). cAMP serves as secondary messenger that activates PKA, leading to the phosphorylation of target proteins and the initiation of downstream events[28]. To investigate whether the regulation of ER Ca^2+^ levels depends on the cAMP-PKA pathway, we employed various methods to elevate basal cytosolic cAMP levels and monitored the corresponding alterations in ER Ca^2+^ levels. We utilized the recently-developed cAMP indicator, cAMPinG1, to monitor basal cAMP levels[29]. Specifically, cells treated with the ACs activator forskolin (FSK) at 0.5 μM showed an approximately 3-fold increase in resting cAMP levels, while 25 μM FSK resulted in an even greater increase of cAMP by approximately 8-fold. Additionally, we examined the effect of overexpressing three different AC subtypes, namely AC2, AC3, and AC9. AC2 and AC9 led to 3.8- and 2.5-fold fold increase, respectively, in the resting cAMP levels; while AC3 did not significantly alter cAMP resting levels (**Fig. 3A**). These interventions produced varied effects on resting ER Ca^2+^ levels. Treatment with 0.5 μM FSK led to a slight increase in ER Ca^2+^ levels (∼1.1-fold), while treatment with 25 μM FSK resulted in a substantial decrease in ER Ca^2+^ levels (∼20%). Interestingly, AC2 did not significantly change ER Ca^2+^ levels, whereas AC3 and AC9 overexpression caused a reduction in ER Ca^2+^ levels by 20% and 30%, respectively (**Fig. 3A**). These results strongly suggest that there seems to be no correlation between cAMP and ER Ca^2+^ levels. The intricate relationship between the dynamics of cAMP levels and ER Ca^2+^ levels implies a complexity that extends beyond a simple linear regulatory relationship. This implies that AC9 might modulate ER Ca^2+^ levels independent of cAMP signaling.

**Figure 3.**
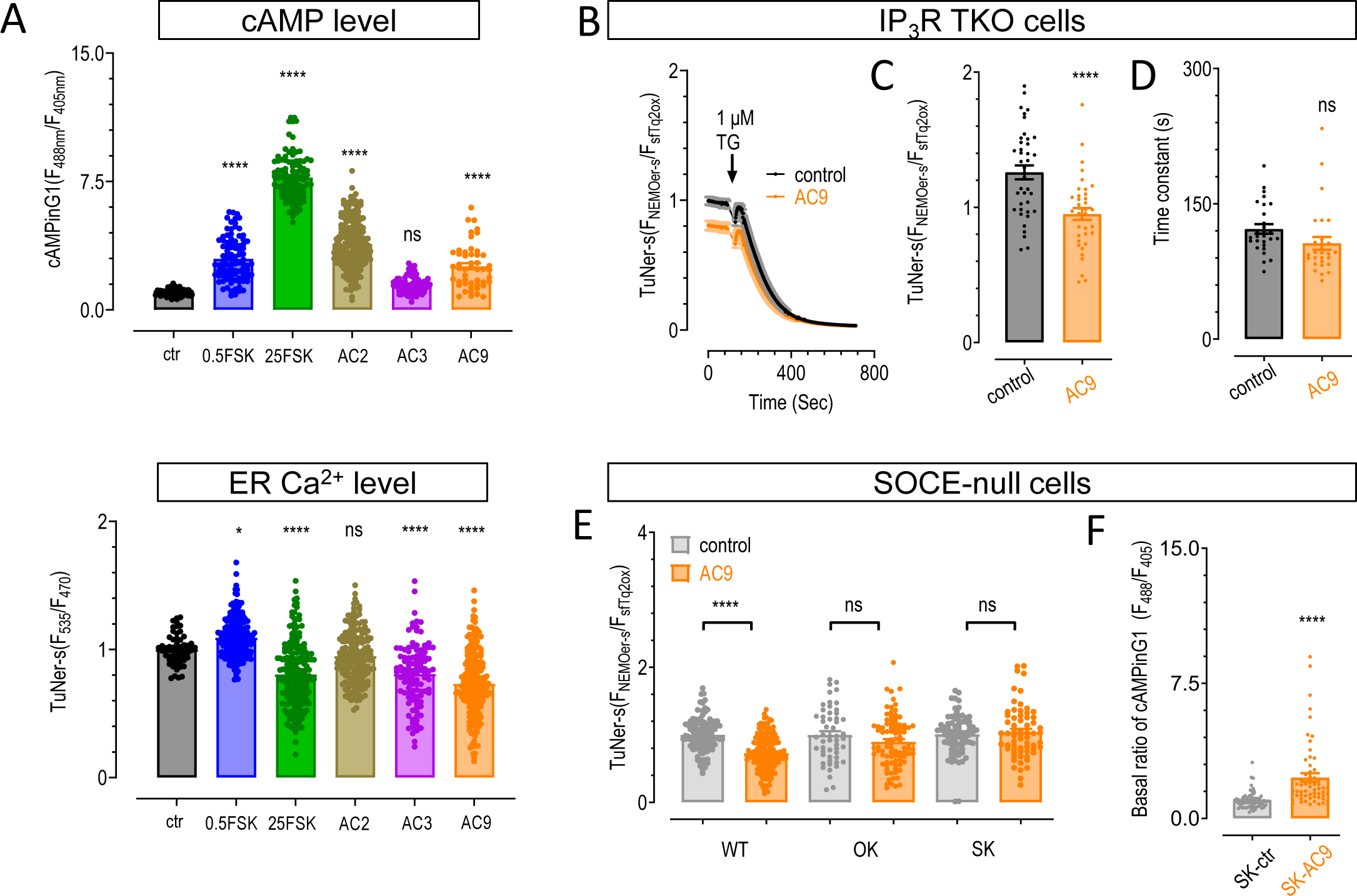
Store operated calcium entry (SOCE), but not IP_3_Rs or cAMP levels, is essential for the AC9-mediated attenuation of ER Ca^2+^ level. A) Effects of activation or overexpression of AC variants on the basal cAMP levels or ER Ca^2+^ levels in HEK293 cells. cAMP levels were presented by the ratio of a cAMP sensor, cAMPinG1 (top), with the ratio of cAMPinG1 normalized to those of the control cells (black: control, n=81 cells; blue: 0.5 μM FSK, n=95 cells; green: 25 μM FSK, n=102 cells; brown: AC2, n=193 cells; purple: AC3, n=85 cells; orange: AC9, n=49 cells. \*\*\*\**P* < 0.0001, ns, not significant, One-way ANOVA) and resting ER Ca^2+^ levels indicated by TuNer-s ratio (bottom). (black: control, n=62 cells; blue: 0.5 μM FSK, n=251 cells; green: 25 μM FSK, n=202 cells; brown: AC2, n=173 cells; purple: AC3, n=111 cells; orange: AC9, n=210 cells. \**P* < 0.05, \*\*\*\**P* < 0.0001, ns, not significant, One-way ANOVA). B-D) In IP_3_R-triple-knock-out (TKO) cells stably expressing TuNer-s (TIRK cells), AC9 still led to a decrease in the resting ER Ca^2+^ level as indicated by TuNer-s signal. Typical traces (B). Statistics of resting TuNer-s ratios (C). Statistics of TG-induced decay time constants of the TuNer-s traces (D). n=39 cells for control and n=37 cells for AC9 overexpression. \*\*\*\**P* < 0.0001, ns, not significant, unpaired Student’s t-test. E) Statistics showing the effects of AC9 on basal ER Ca^2+^ levels in the wild type (WT), STIM1/2-double Knockout (SK) cells and Orai1/2/3-triple Knockout (OK) HEK293 cells (For WT cells, n=111 cells in control group and n=179 cells in AC9 group; For OK cells, n=53 cells in control group and n=91 cells in AC9 group; for SK cells, n=64 cells in control group and n=81 cells in the AC9 group. \*\*\*\**P* < 0.0001, unpaired Student’s t-test). F) Statistics showing the effects of AC9 on cAMP levels in the SK cells (n=74 cells for control and n=60 cells for the AC9 group. \*\*\*\**P* < 0.0001, ns, not significant, unpaired Student’s t-test). Data are shown as mean ± SEM of at least three independent experiments.

We next examined whether AC9 affects ER Ca^2+^ levels by influencing IP_3_Rs, key players for the maintenance of ER Ca^2+^ homeostasis. We overexpressed AC9 in <u>T</u>uNer-s expressing cells with IP_3_<u>R</u>1/2/3 triple-<u>k</u>nockout (TIRK cells)[30] and assessed ER Ca^2+^ levels before and after TG-induced Ca^2+^ store depletion. We found that overexpression of AC9 still significantly lowered ER Ca^2+^ levels in TIRK cells (**Fig. 3B-C**), suggesting that AC9 could perturb ER Ca^2+^ homeostasis independent on IP_3_R expression. Furthermore, the rate of TG-induced Ca^2+^ decay remained unaltered, indicating that AC9 had no appreciable effect on ER Ca^2+^ leak channels (**Fig. 3D**).

### AC9 reduces ER Ca^2+^ levels by suppressing store-operated Ca^2+^ entry (SOCE)

Next, we moved on to test whether AC9 could affect ER Ca^2+^ levels through modulating the SOCE pathway. We utilized Orai 1/2/3 triple-knockout (OK) cells or STIM1/2 double-knockout (SK) cells[31] devoid of key components of SOCE. Compared to SK or OK cells expressing TuNer-s alone, SOCE-null cells co-expressing TuNer-s and AC9 showed a similar basal TuNer-s ratio (**Fig. 3E**). This finding indicates that AC9 was no longer able to lower ER Ca^2+^ levels as seen in wild-type cells tested side-by-side (**Fig. 3E**). In parallel, we measured the basal cAMP levels with cAMPinG1 in SK cells transiently co-expressing AC9. We detected an increase in cAMP levels, even though AC9 failed to cause a reduction in ER Ca^2+^ levels in these cells (**Fig. 3F**). Clearly, AC-catalyzed cAMP production does not seem to be implicated in lowering ER Ca^2+^ levels. Instead, SOCE is found to be essential for AC9-mediated attenuation of ER Ca^2+^ content.

We subsequently explored the effects of AC9 on TG-induced SOCE (**Fig. 4A**). As shown by changes in TurN-s ratio that reports cytosolic Ca^2+^ changes, we noted that AC9 could significantly reduce TG-induced SOCE (**Fig. 4A-B**). We further examined the effect of SOCE inhibition on ER Ca^2+^ content. Consistent with previous reports[21, 32], cells with transient knockdown of STIM1 and diminished SOCE showed reduced basal ER Ca^2+^ levels, as indicated by lower TuNer-s ratio (60 ±1.7 % of control) (**Fig. 4C-D**). Together, these data converge to support the conclusion that AC9 lowers the ER Ca^2+^ content by acting on the SOCE pathway.

**Figure 4.**
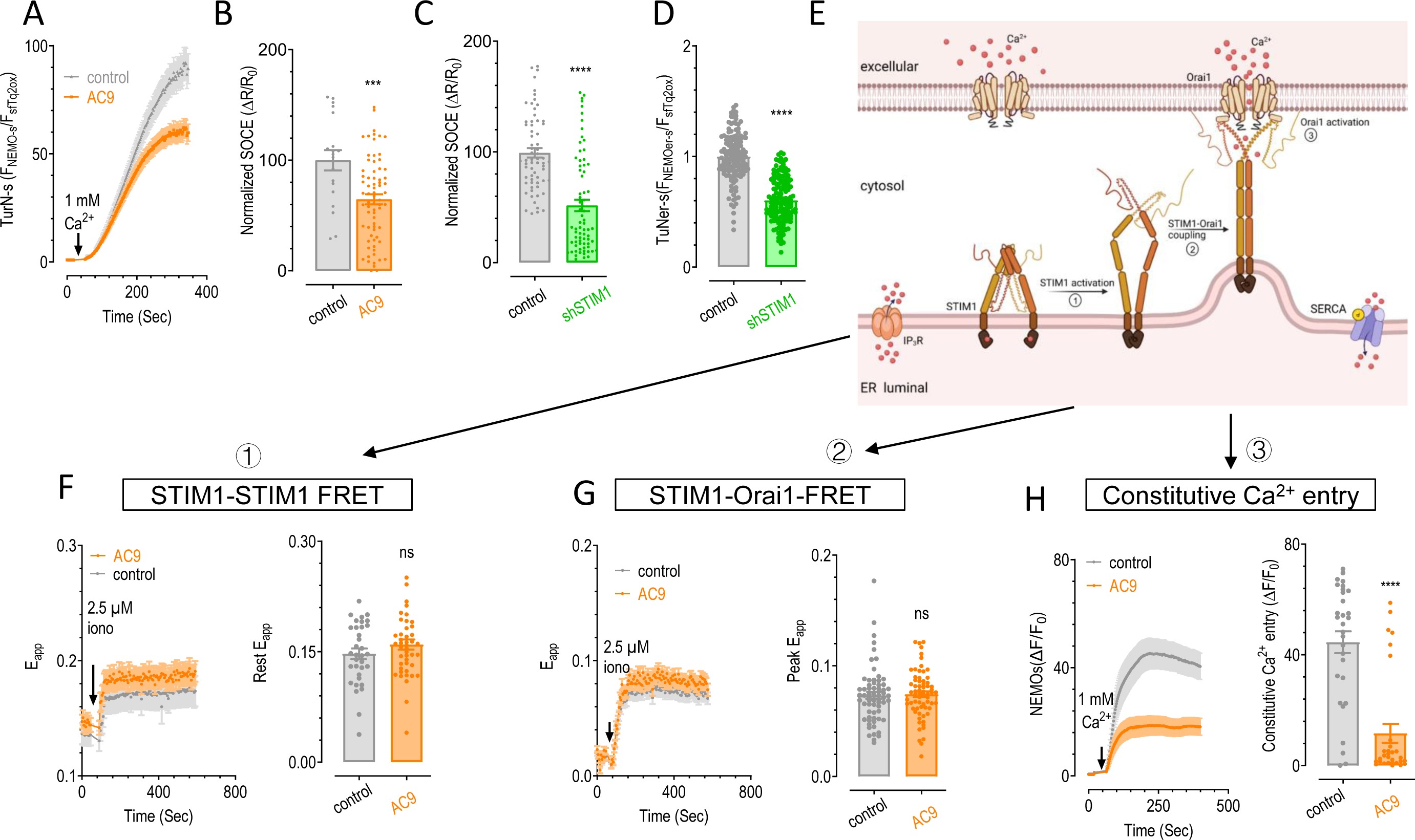
AC9 regulates ER Ca^2+^ levels by inhibiting SOCE via its action on Orai1. A-B) The inhibitory effects of AC9 (orange) on SOCE in HEK 293 cells stably expressing cytosolic Ca^2+^ indicator TurN-s. Prior to recordings, cells were bathed in nominally Ca^2+^ free solution containing 1 μΜ TG for 10 min to empty ER Ca^2+^ stores. Typical traces (A) and statistics of mean SOCE responses normalized to the internal control (B) (n=21 cells for internal control and n = 72 cells for the AC9 group. \*\*\**P* = 0.001, \*\*\*\**P* < 0.0001, unpaired Student’s t-test). C-D) Statistics showing the effects of STIM1 knockdown on SOCE responses (C) and basal ER Ca^2+^ levels (D) (SOCE: n = 65 cells for control and n=73 cells for shSTIM1. ER Ca^2+^ levels: n = 150 cells for both groups. **** *P* < 0.0001, unpaired Student’s t-test). E) Cartoon depicting the key activation process of SOCE, involving three steps: ① Activation of STIM1; ② Coupling of STIM1 with Orai1; ③ Activation of the Orai1 channel, resulting in channel opening and the influx of extracellular Ca^2+^ into the cell. F-H) Effects of transient overexpression of AC9 on key steps of SOCE activation. Left, typical traces; right, statistics. FRET responses between YFP-STIM1 and CFP-STIM1 (F) (n = 34 cells for control and n = 45 cells for the AC9 group. ns, not significant, unpaired Student’s t-test). FRET responses between YFP-STIM1 and CFP-Orai1 (G) (n = 66 cells for control and n=67 cells for the AC9 group. ns, not significant, unpaired Student’s t-test.). The constitutive Ca^2+^ entry in cells transiently expressing full length Orai1-ANSGA (H) (n = 30 cells for control and n=31 cells for the AC9 group. \*\*\*\**P* < 0.0001, unpaired Student’s t-test). Data are shown as the mean±SEM of at least three independent experiments.

### AC9 attenuates SOCE by impeding the function of Orai1

To further investigate the inhibitory mechanism of AC9 on SOCE, we systematically analyzed its effects on key steps in the SOCE activation pathway (**Fig. 4E-H**). We first employed fluorescence resonance energy transfer (FRET) assays to monitor the oligomerization of STIM proteins[33], which represents the initial major step during SOCE activation. The results showed that the FRET signals between YFP-STIM1 and CFP-STIM1 remained largely unaltered in the presence of AC9, irrespective of the resting state or after treatment with 2.5 μM ionomycin to induce store depletion (**Fig. 4F**). Therefore, the initial activation of STIM1 during SOCE seems to be unaffected by AC9.

The second major step in SOCE activation involves the translocation of STIM1 toward ER-PM junctions, where it couples with Orai1 through physical interactions (**Fig. 4E**). We therefore examined whether the expression of AC9 could influence this step by monitoring the FRET signal between STIM1-YFP and Orai1-CFP[34]. Our results revealed that overexpression of AC9 neither altered the resting FRET signal between STIM1-YFP and CFP-Orai1, nor the FRET signal after ionomycin-induced SOCE activation (**Fig. 4G**). Congruently, this finding suggests that AC9 does not significantly impact the coupling process between STIM1 and Orai1.

The third key step in SOCE activation involves Orai1 undergoing conformational changes upon STIM1 engagement, subsequently causing the opening of Orai1 channel to mediate Ca^2+^ influx (**Fig. 4E**). To assess whether AC9 could directly act on Orai1, we examined its effects on Orai1-ANSGA, an Orai1 mutant that remains constitutively active in the absence of STIM1 binding, in HEK293 cells stably expressing the ultra-sensitive Ca^2+^ indicator NEMOs (termed as NEMOs cells)[35]. Compared with cells expressing Orai1-ANSGA alone, those co-expressing Orai1-ANSGA and AC9 reduced the cytosolic Ca^2+^ levels, as reflected by lower NEMO-s fluorescence intensities (**Fig. 4H**). Our observations revealed a significant inhibition of constitutive Ca^2+^ influx by AC9, indicating its suppressive effect on Orai1 channels *per se*.

To further pinpoint the specific regions within Orai1 that are important for the suppressive effect imposed by AC9, we generated several truncated versions of Orai1-ANSGA, namely Orai1-ANSGA_65-301_ with the N-terminus truncated, Orai1-ANSGA_1-266_ with the C-terminus truncated, and Orai1-ANSGA_65-266_ with the N/C termini removed. Worthy to note, NEMOs cells overexpressing any of the three mutants still exhibited constitutive Ca^2+^ influx, indicating the functional integrity of these variants (**Fig. 5A-C**). Reduced Ca^2+^ entry mediated by Orai1-ANSGA1-266 may be attributed to its lower expression. Co-expression of AC9 significantly inhibited the spontaneous Ca^2+^ influx mediated by Orai1-ANSGA_65-301_ (**Fig. 5A**), while it did not seem to alter the constitutive Ca^2+^ influx mediated by Orai1-ANSGA_1-266_ or Orai1-ANSGA_65-266_ (**Fig. 5B-C**). Together, these results indicate that the C-terminus of Orai1 is required for AC9 to exert its inhibitory effect on Orai1. Interestingly, even though it is well established that the C-terminus of Orai1 is essential for the binding of STIM1[11, 36], we found no detectable interference of AC9 on the functional coupling between STIM1 and Orai1, as indicated by unaltered FRET signals (**Fig. 4G**). It is plausible that, apart from interacting with STIM1, the C-terminus of Orai1 is subject to regulation by alternative factors such as AC9, an intriguing concept that merits detailed exploration in the near future.

**Figure 5.**
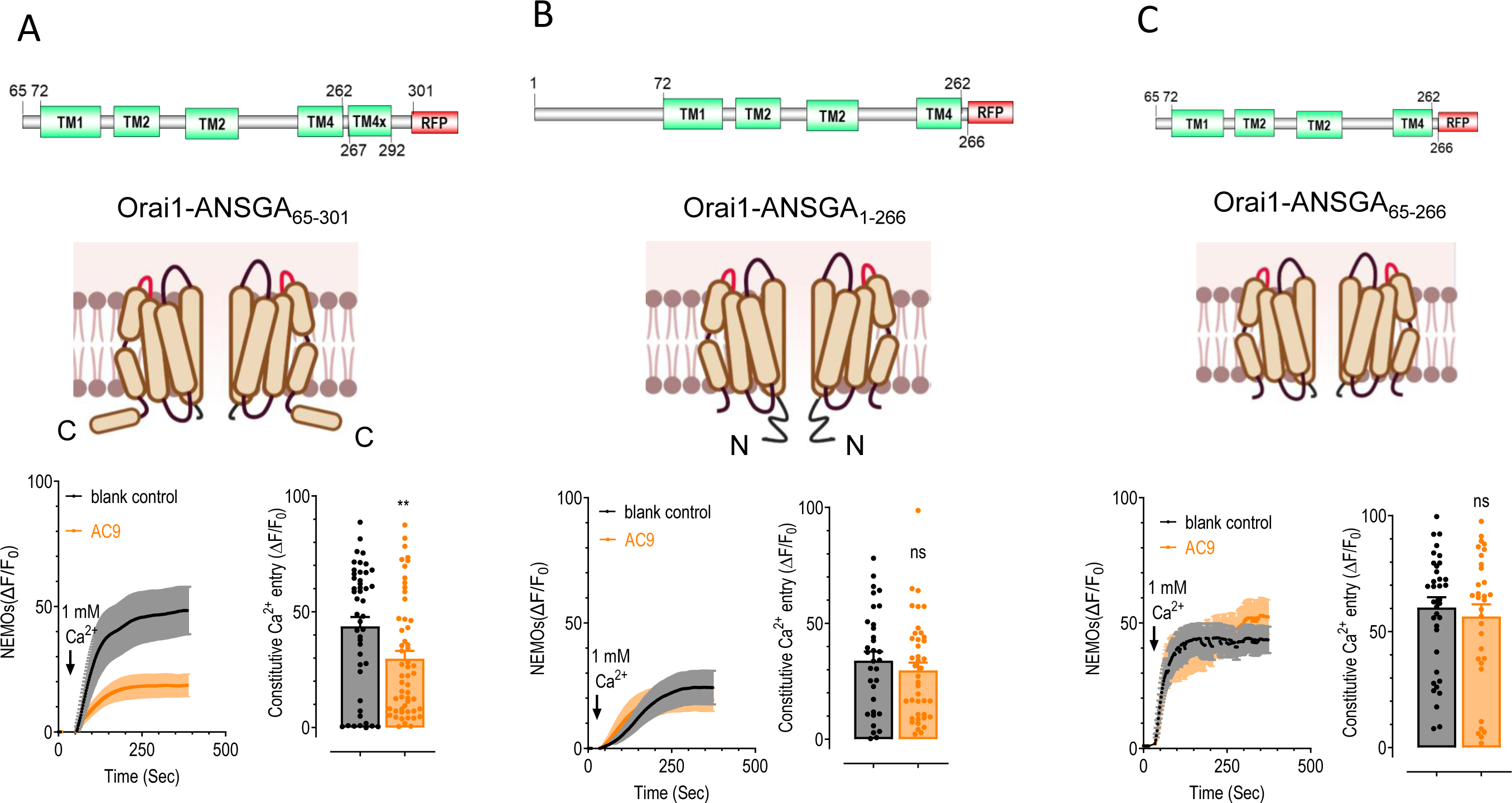
AC9 inhibits the function of Orai1 by acting on Orai1 C-terminus. Effects of AC9 on the constitutive Ca^2+^ entry caused by transiently expressing truncated Orai1-ANSGA variants in HEK293 cells. The cytosolic constitutive Ca^2+^ entry level was monitored by Ca^2+^ indicator NEMOs. Cartoon illustration of tested Orai1-ANSGA variants (TM: transmembrane domain; TM4x: cytosolic e<u>x</u>tension of TM4) (top). Typical traces (bottom left). Statistics of plateau constitutive Ca^2+^ entry (bottom right). Cells transiently expressing Orai-ANSGA with the N terminus deleted (ANSGA_65-301_) (A); deletion of the C terminus, ANSGA_1-266_ (B) and removal of both termini, ANSGA_65-266_ (C) (ANSGA_1-266_, n = 33 cells; ANSGA_1-266_+AC9, n = 41 cells; ANSGA_65-301_, n = 46 cells; ANSGA-65-301+AC9, n=55 cells; ANSGA-65-266, n = 37 cells; ANSGA-65-266+AC9, n = 33 cells. \*\**P*<0.01; ns, not significant, unpaired Student’s t-test). Data are shown as mean ±SEM of at least three independent experiments.

### AC9 is involved in lowering ER Ca^2+^ levels during *Drosophila* brain development

To explore whether AC9 is a conserved key factor regulating the decrease in ER Ca^2+^ levels during neuronal development *in vivo*, we went on to examine its expression and functional roles in *Drosophila* brain neurons. We first investigated the expression pattern of AC9 during the development of *Drosophila*. We selected third-instar larvae and adult flies 3 days post-eclosion and compared the expression of Ac13E (an ortholog of human AC9). In line with observations made in the rodent brain (**Fig. 2F**), the mRNA level of Ac13E in the adult fly brain was upregulated by 31.8-fold compared to the larval brain (**Fig. 6A**). These results demonstrate that AC9 undergoes upregulation during the developmental processes of both *Drosophila* and rodents, indicating its conservation across these species.

**Figure 6.**
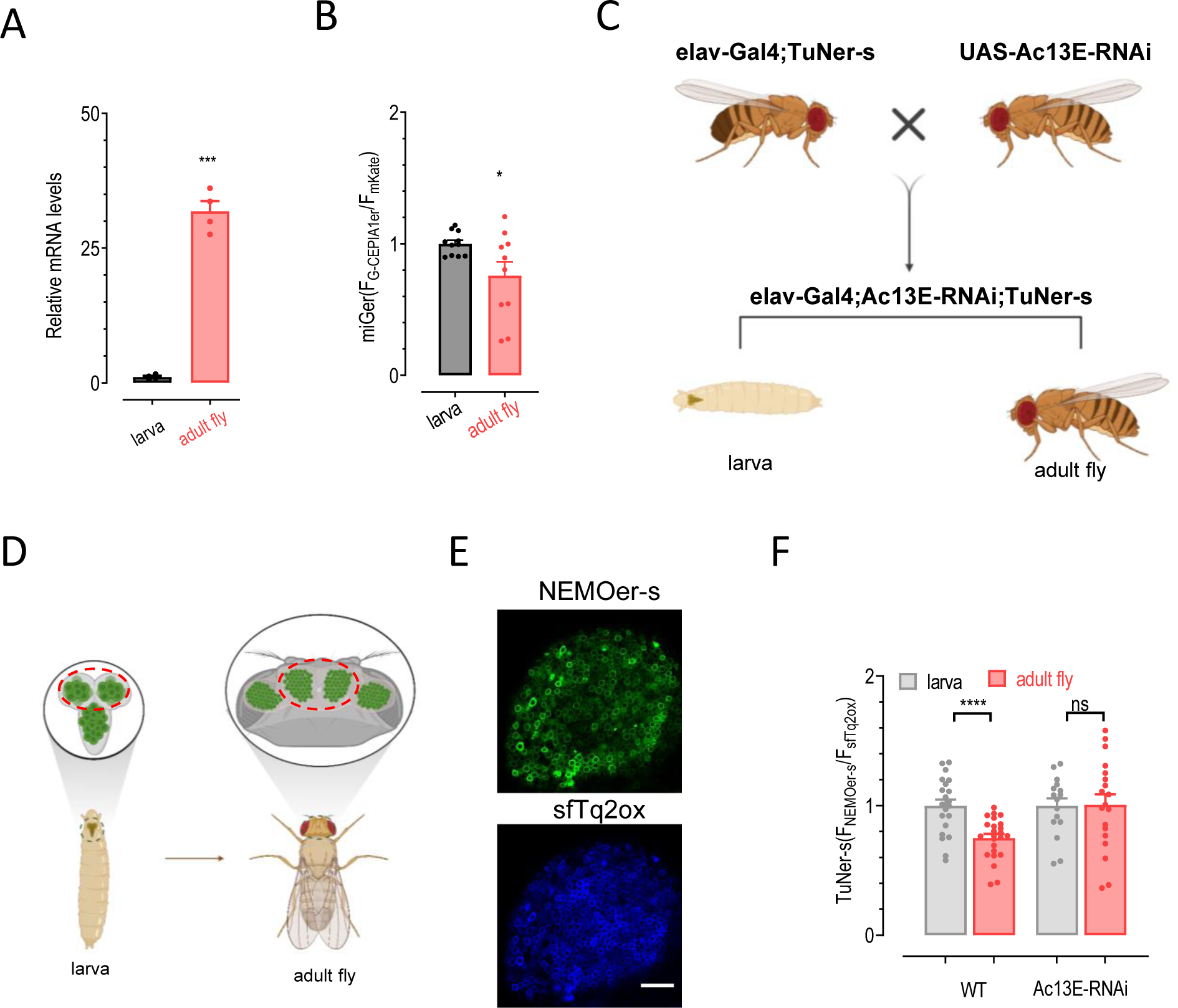
Upregulation of AC9 in neurons is essential for reducing ER Ca^2+^ levels during *Drosophila* brain development. A) Relative mRNA levels of AC9 in brains of larval or adult *Drosophila*. The mRNA level of RPL32 was used as an internal control (Data are shown as the mean ±SEM of four independent experiments, each from the average of at least 3 technical replicates. \*\*\**P* < 0.001. paired Student’s t-test). B) Statistics of ER Ca^2+^ levels in neurons derived from larval and adult brains of *Drosophila* indicated by miGer. The miGer ratios of adult fly brains were normalized to the ratios of larval brains (n=11 for larva and n=10 for adult fly. \**P* < 0.05. unpaired Student’s t-test). C) Schematic of a genetic cross to generate the fly strains expressing TuNer-s, with Ac13E (the homology gene of AC9 in *Drosophila*) knocked down in neurons driven by elav-Gal4. D) Illustration of the microscopic view of brain from the larval (left) and adult (right) *Drosophila* with neuron-specific expression of TuNer-s. Red dash circles represent the regions of recording. E) Typical fluorescence images showing TuNer-s signals in the central brain regions of adult *Drosophila* monitored with confocal imaging. The fluorescence of NEMOer-s (top). The fluorescence of sfTq2ox (bottom). Scale bar, 10 μm. F) Statistics showing the basal ER Ca^2+^ levels in neurons of WT or Ac13E-KD *Drosophila* larval or adult brains, which is indicated by the TuNer-s ratios. The TuNer-s ratios of adult fly brains were normalized to the ratios of larval brains (n = 21, WT larval and n = 24, WT adult, n = 16 for kd-AC9 larval and n=19 for kd-AC9 adult, each representing the average of 30 cells from each brain. \**P* < 0.05, ns, not significant, unpaired Student’s t-test).

We next sought to monitor changes in ER Ca^2+^ levels in *Drosophila* brain neurons during development. To achieve this, we generated a transgenic fly strain with pan-neuronal expression of miGer, driven by the elav-Gal4 promoter. Subsequently, we examined the miGer ratios in neurons within the central brain region of third-instar larvae and adult flies three days post-eclosion. We observed a significant reduction in miGer signals in the adult fly brains compared to larvae (**Fig. 6B**). This observation is consistent with previous reports showing reduced ER Ca^2+^ release in neurons of various animal species[37, 38]. To the best of our knowledge, this represents the first direct measurement of ER Ca^2+^ content during development.

To explore whether Ac13E is essential for the decrease in ER Ca^2+^ levels during the developmental process of *Drosophila* brain neurons, we first generated a fly strain expressing the more sensitive ER Ca^2+^ indicator, TuNer-s, promoted by elav-Gal4 driver. Subsequently, we crossed this strain with Ac13E knockdown (KD) flies to generate TuNer-s Ac13E KD flies (**Fig. 6C-D**). The generated strain exhibited concurrent knockdown of Ac13E and overexpression of TuNer-s in neurons (**Fig. 6E**). Larvae and adult fly brains from this strain were selected to measure ER Ca^2+^ levels in neurons. The results indicated that, upon Ac13E knockdown, there was no significant reduction in ER Ca^2+^ levels observed between larval and adult fly neurons (**Fig. 6G**). Collectively, these results firmly validate that AC13E plays a crucial role in lowering ER Ca^2+^ levels during the developmental process of *Drosophila*.

### Conclusion

In summary, by employing a genome-wide editing approach and Ca^2+^ imaging analysis, we have identified AC9 as a critical regulator of ER Ca^2+^ homeostasis, the overexpression of which leads to a reduced ER Ca^2+^ level. At the mechanistic level, AC9 seems to act on Orai1 to inhibit SOCE, ultimately causing a decline in the ER Ca^2+^ content. Physiologically, the increased expression of AC9 is associated with the reduction of ER Ca^2+^ levels during brain development. Therefore, our study provides novel insights into the biological role of AC9 in regulating ER Ca^2+^ homeostasis and developmental processes.

## Methods

### Plasmid construction

To generate the mScarlet-tagged candidate construct, the coding sequences (CDS) of the candidate genes were PCR-amplified from the human cDNA library (kind gift from Professor Qian Zhaohui) or obtained from the Bio-Research Innovation Center Suzhou. The mScarlet CDS sequence was PCR-amplified from mScarlet-MYO10 (addgene Plasmid #145179) and subsequently subcloned into the pcDNA3.1(+) backbone using a multiple-fragment homologous recombination kit (Cat#: C113, Vazyme biotech, Nanjing, China).

For the construction of Cas9-p2A-miGer, the Cas9 and miGer sequences were PCR-amplified from lentiCas9 and pCDNA3.1-miGer, respectively. Subsequently, these sequences were ligated into pcDNA3.1(+) using primers that incorporated the P2A sequence (5′-TCTAGAGGTGGCGGCTCAGGATCC-3′) between the EcoRⅠ and EcoRⅤ sites. To construct the AC9 knockdown plasmid, we design the gRNA targeting the CDS of AC9 as follows: 5’-ACCTCCAGTCCCAAGAACAGGA-3’. The gRNA oligos were annealed and inserted into the CasRx vector (Addgene: # 134842) linearized by BsmBI (New England Biolabs) with T4 ligase (New England Biolabs). To generate the STIM1 knockdown plasmid, the shRNA sequences used to target STIM1 were as follows: 5’-CCTGGATGATGTAGATCATAA-3’, inserted to the pLKO.1 vector between the AgeⅠ and EcoRⅠ sites.

### Cell culture and transfection

HEK 293 cells were cultured in Dulbecco’s modified Eagles’ medium (DMEM) supplemented with 10% fetal bovine serum (FBS), 1% penicillin/streptomycin, and maintained at 37°C in a humidified environment with 5% CO2. Mouse N2a cells were cultured in MEM supplemented with 10% fetal bovine serum, and 1% penicillin/streptomycin. To induce N2a cell differentiation, MEM containing 0.1% FBS and 20 μM retinal acid was used to treat cells for at least 3 days.

For HEK 293 cells, transfections were performed by electroporation using the Bio-Rad Gene Pulser Xcell system (Bio-Rad, Hercules, CA, USA) in 4 mm cuvettes and OPTI-MEM medium-and a voltage step pulse (180 V, 25 ms, in 0.4 ml of the medium) was used. For N2a cells, transfections were performed with purified plasmids using lipofectamine 3000 (Invitrogen, Waltham, MA, USA) according to the manufacturer’s instructions.

To establish stable cells, the Ca^2+^ indicator TuNer-s was transfected into HEK293 cells, N2a cells, and IP_3_R triple-knockout cells. After selection with 2 μg / ml puromycin for 5-7 days, the cells were then diluted to single clones and expanded in culture. Healthy clones with high expression and normal Ca^2+^ responses were selected for usage. Cas-p2A-miGer stable cells were established using the same process.

### Genome-wide CRISPR-Cas9 screening

Well-established genome-scale CRISPR knockout (GeCKOv2) library[39] targeting 19050 annotated genes with 123411 sgRNAs were transduced to HEK 293 cells stably expressing Cas9-p2A-miGer with a multiplicity of infection (MOI) of five. The infected cells were selected by puromycin for 5 days to construct genome-edited cell pools. Cells were then sorted based on miGer ratios using flow cytometry. Compared with control group, cells with elevated ratio and those with reduced ratio were collected and cultured. When the sorted cells expanded to 3×10^7^, they were collected to extract genomic DNA, and amplified the sgRNA for next-generation sequencing (NGS) performed by commercial vendor (syngentech). We conducted gene enrichment analysis using clusterProfiler (version 4.8.1) and annotated the results with the org.Hs.eg.db package (version 3.17.0). Differential expression analysis was performed Qlucore Omics Explorer (version 3.4).

### Western blotting

The lysis of Cas9-p2A-miGer cells was performed using cell lysis buffer (BestBio). After 30 minutes of lysis, the sample was centrifuged, and the supernatant was collected. Protein concentration was quantified using the Bradford protein assay. Aliquots of 20 µg of protein were prepared and subjected to SDS-PAGE electrophoresis. Subsequently, they were transferred to polyvinyl difluoride membranes (Millipore). The membranes were blocked with 5% skim milk at 4 °C overnight, followed by incubation with mouse Anti-Flag primary antibody (1:500) for 2h and membranes were washed 3 times (10 min) in PBST and incubated with goat anti-mouse secondary antibody conjugated with horseradish peroxidase for 2h. Internal control GAPDH was detected with anti-GAPDH antibody (sc-32233; SANTA) (1:2000 dilution) and corresponding secondary antibody is anti-mouse-IgG (7076S; CST) (1:2000 dilution). Subsequently, membranes were washed 3 times (10 min) in PBST. Detection was examined with chemiluminescent substrate (Millipore) using the manufacturer’s protocols.

### Fly stocks, rearing, and brain dissection

UAS-miGer (attp2) and UAS-TuNer-s (attp2) flies were generated by Fungene Biotechnology. To achieve neuron-specific expression, they were crossed with the flies containing the elav-GAL4^c155^ driver. UAS-Ac13E RNAi Drosophila were obtained from Bloomington Drosophila Stock Center (62247). All strains were grown in standard corn flour agar incubators at 25 °C.

Dissection of larval and adult brains was carried out as previously described[40]. For Larval brain dissection, we used forceps to rip and remove the larval cuticle, severed the brain from the midgut, and removed fat tissue and imaginal discs. For adult brain dissection, we first cut off the head, next peeled away the cuticle, and then removed the trachea until the brain is isolated. During dissection, flies were immersed in PBS, and dissected brains were kept in Ca^2+^ imaging solution for imaging experiments or in lysis solution for RNA extraction.

### Animals and tissue collection

This study included two separate animal experiments: Neonatal (1-3 days) and adult Sprague Dawley rats (male; 6-8 weeks; 200-250g) were derived from Viton Lihua Corporation (Beijing, China). All adult rats were placed in clear, polyethylene cages with a relative humidity of 40-60% and maintained under standard laboratory conditions (22-25℃), with a 12-h light/dark cycle after 1-week acclimatization. Standard laboratory food and tap water were available ad libitum. All adult rats were fasted and refrained from water overnight before the experiments. All experiments were approved in accordance with the Animal Care and Ethics Committee.

Neonatal rats were sacrificed through cervical dislocation. Adult rats were sacrificed by decapitation after peritoneal anesthesia with pentobarbital sodium (50 mg·kg^-1^). After alcohol disinfection, brain tissue samples were gently extracted from the cranial cavity, utilizing the foramen magnum as the lower limit. The samples were then dissected free of meninges and superficial blood vessels immediately on the ice table. The cerebral cortexes were carefully dissected from the brains on an ice table and placed in frozen tubes, stored in a liquid nitrogen tank.

### RNA isolation, RT–PCR, and quantitative PCR (qPCR)

Total RNA was extracted from cells, rat cerebral cortexes or fly brains using TRIzol and the reverse transcription was performed with PrimeScript™ RT Master Mix (Takara) following the manufacturer’s instructions. The cDNA product was used as a template for qPCR run and mixed with primers and SYBR Green PCR Master Mix. qPCR reactions was performed on QuantStudio™ 6 Flex Real-Time PCR System. Relative mRNA levels were calculated using Comparative Ct (△△CT) method. The data from rat samples were normalized to corresponding GAPDH levels, and from *Drosophila* samples were normalized to RPL32 levels. Primer sequences used were listed in the following table.

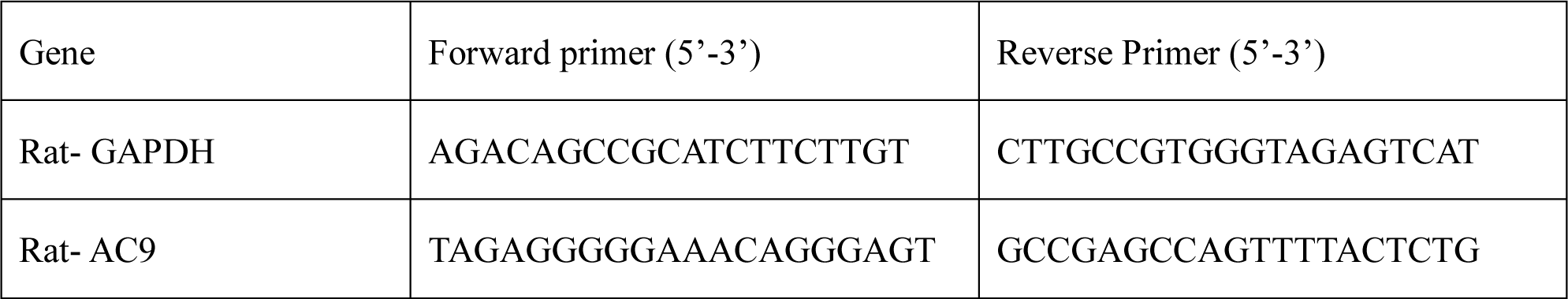
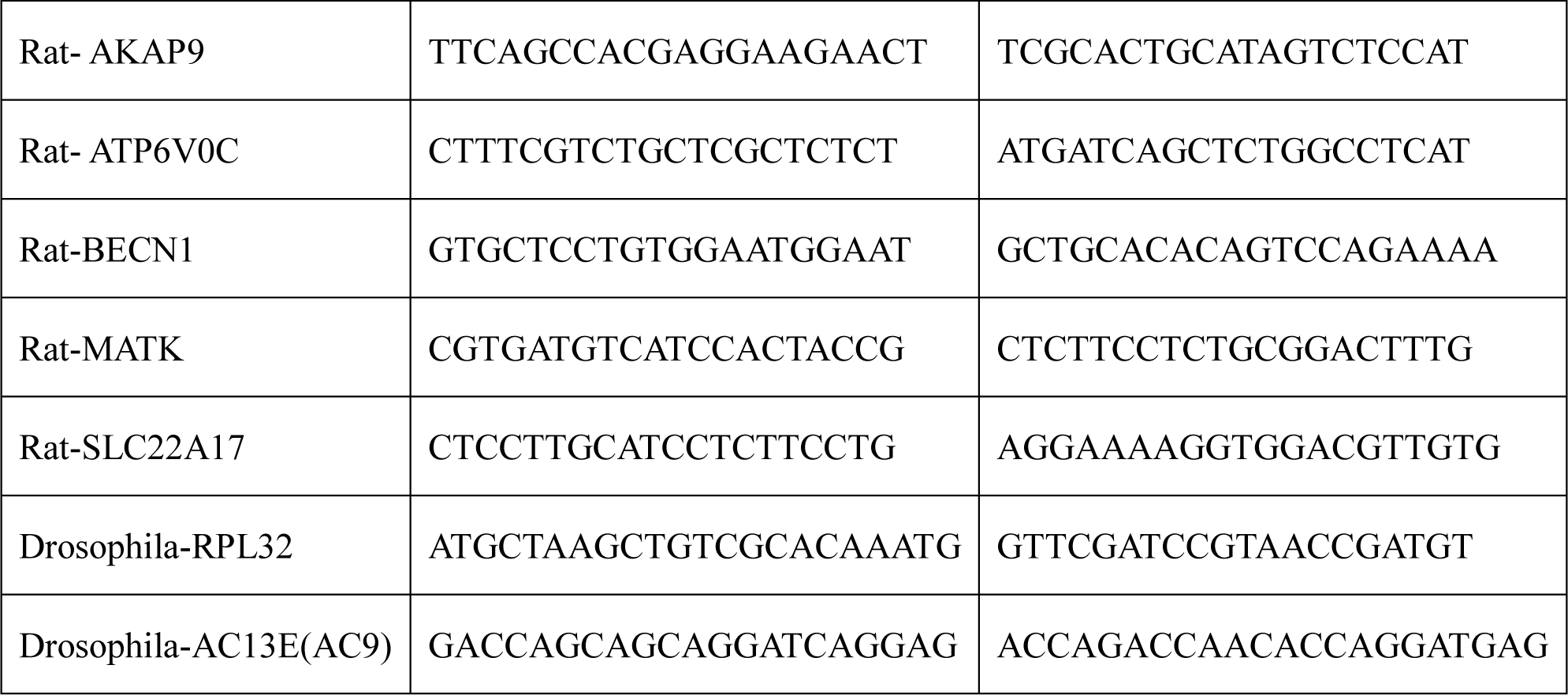

### Live-cell fluorescence imaging

Cells seeded on the round coverslips were transferred to an imaging chamber and incubated in imaging buffer which containing 107 mM NaCl, 7.2 mM KCl, 1.2 mM MgCl_2_, 11.5 mM glucose, and 20 mM HEPES-NaOH (pH 7.2). Fluorescence signals were acquired with a ZEISS observer Z1 imaging system controlled with the SlideBook software v.6.0.23 (Intelligent Imaging Innovations, Inc.). Ca^2+^ signals were indicated by ratiometric-GECIs: miGer, TuNer-s or TurN-s and monomeric ER Ca^2+^ indicator NEMOs. Filters for miGer (mKate: 549 ± 6 nm _Ex_, 628 ± 48 nm _Em_; G-CEPIA1er: 470 ± 11 nm _Ex_, 510 ± 14 nm _Em_), and for TuNer-s or TurN-s (NEMOs or NEMOer-s: 500 ± 10 nm _Ex_; 535 ± 15 nm _Em_; sfTq2^OX^: 438± 12 nm _Ex_, 470 ± 12 nm _Em_) were used. Ca^2+^ levels are indicated as F_G-CEPIA1er_/F_mkate_ ratio or F_NEMOs_/F_sfTq2OX_ ratio after background subtracted. Images were captured every 2 s. FRET measurements were performed with the CFP (438 ±12 nm _Ex_; 470 ± 12 nm _Em_), YFP (500 ± 10 nm _Ex_; 535 ± 15 nm _Em_), and FRET_raw_ (438 ± 12 nm _Ex_; 535 ± 15 nm _Em_) filters, capturing images every 10 s. Apparent FRET efficiency (Eapp) values were calculated using the same method as those previously described[41]. cAMP level was indicated by ratiometric indicator cAMPinG1 (377 ± 25 nm _Ex_ and 470 ± 12 nm _Ex_; 525 ± 25 nm _Em_). The corresponding fluorescence readings from regions of interest (ROI) were exported from the software and imported into MATLAB 2014a (The MathWorks, Natick, MA, USA) and plotted with GraphPad Prism 9.51 software. The ratio images were generated and assigned pseudocolors using Slidebook software (v.6.0.23). All experiments were carried out at room temperature. Traces shown are representative of at least three independent experiments with each including at least 30 cells.

### *Drosophila* brain ex vivo imaging

For drosophila brain imaging, dissected brains were mounted into the coverslips coated with poly-L-lysine and immersed in an imaging buffer containing 1mM Ca^2+^. The signals were collected by the ZEISS LSM880 microscope equipped with a 63x oil objective (NA 1.4). miGer (G-CEPIA1er/ mKate) was excited by 488 nm and 543 nm laser respectively and the resulting fluorescence at 490 - 590 nm and 590 - 690 nm was collected. TuNer-s (NEMOer-s/sfTq2^OX^) was excited by 514 and 405 nm laser respectively and detected at 560 - 650 nm and 420 - 550 nm. The signals of ROIs were analyzed using FIJI software and plotted with Prism 9.51 software.

### Long-term Ca^2+^ imaging

N2a cells stably expressing miGer were seeded on glass-bottom 24-well plates (Cellvis, Cat#: p24-1.5H-N) and cultured in MEM supplemented with 2% FBS and 40 μM retinal acid to induce differentiation. Long-term imaging was then performed by using the Livecyte^TM^ system (Phase Focus) with an acquisition interval of 2h for 3 days. Cells were tracked automatically with Livecyte Analyze software. The fluorescence signals of each cell were exported as txt files and plotted with Prism7 software.

### Statistical analysis

All quantitative data are presented as means ± SEM of at least three independent biological repeats. Comparisons between the two groups were analyzed via unpaired t-test. Comparisons among multiple groups were analyzed with One-way ANOVA.

## Author Contributions

Y. W. conceived the project, designed the experiments. Y. W. and L. W. wrote the manuscript. J. L. performed CRISPR-screen. YP.W. and Z.Z. did bioinformatics analysis. L.W., YP. W., Z.Z., YQ. W., R.H generated candidate-mScarlet plasmids and Ca^2+^ imaging assays. L. W. performed all the other experiments and did data analysis. P.L. collected rat cerebral cortexes. E.Y. and R.C. helped with the generation of TuNer-s stable *drosophila* strain.

## Competing Interests

The authors have declared no competing interests.

## Acknowledgements

This work was supported by the Ministry of Science and Technology of China (2019YFA0802104 to Y.W.), the National Natural Science Foundation of China (92254301and 91954205 to Y.W.).

## Data Availability Statement

The data that support the findings of this study are available upon reasonable request.

